# Nanoparticles reveal permanent and reversible changes to lymph node biomechanics during inflammatory response

**DOI:** 10.1101/2025.02.24.639945

**Authors:** Ann Ramirez, Vedanth Sriram, Yassmin Abbouchi, Reina Patolia, Emily Passaro, Michele Kaluzienski, Katharina Maisel

## Abstract

Lymph nodes are highly specialized immune organs that orchestrate the adaptive immune response. In the lymph nodes, naïve B and T lymphocytes encounter cognate antigens, sparking their activation and response to foreign substances. Lymph nodes grow in response to an immune challenge, at least in part to accommodate increased numbers of infiltrating and proliferating B and T lymphocytes. This behavior is supported by a robust three-dimensional network of extracellular matrix (ECM) fibers and fibroblastic reticular cells (FRCs). ECM fibers and FRCs work synergistically to alternate stretching and contractile forces between them allowing the lymph node to maintain structural integrity during rapid tissue reconstruction. These changes ultimately alter the material properties of the lymph node, which can impact cell migration, proliferation, and differentiation. Recent work has investigated the physiological implications of the changing lymph node microenvironment; however, the biophysical properties of the lymph nodes during these changes remain largely unexplored. Here, we use multiple particle tracking microrheology (MPT), a minimally invasive nanoparticle-based technique to investigate the biophysical properties (elastic/loss moduli, microviscosity, pore size) of lymph nodes post inflammatory stimulus. Our results highlight mechanical changes both during the initial phases of the acute inflammatory response and upon resolution of inflammation, a topic that is relatively understudied. We show that B and T cell rich areas exhibit comparable changes in biomechanical properties over time, suggesting that they restructure in a similar fashion during acute inflammation.

Additionally, for the first time, we show that biological sex modulates lymph node biomechanics in acute inflammation: Lymph nodes from female mice showed a ∼20-fold increase in elastic and loss moduli at peak inflammation, while lymph nodes from male mice had a ∼5-fold decrease in both moduli. Additionally, lymph nodes from female mice appeared to permanently remodel during the resolution of acute inflammation resulting in the maintenance of an overall higher elastic and loss modulus, while lymph nodes from male mice returned to the biomechanics of untreated lymph nodes. We also found that at least some of the changes in biomechanical properties were correlated with changes in ECM materials in the lymph nodes, suggesting a structure-function relationship. Overall, our studies provide key insights into how biomechanical properties in lymph nodes are altered during inflammation, a previously unstudied area, and lay the foundation for structure-function relationships involved in immune response. Additionally, we demonstrate a robust technique for the analysis of the lymph node interstitial tissue properties and how they vary with inflammatory stimuli.

## INTRODUCTION

The lymph node is key in shaping the adaptive immune response. The complex coordination of adaptive immune cell (lymphocyte)-mediated immune responses is largely due to the lymph node’s highly organized structure. The lymph node can be separated into three distinct regions: the cortex, paracortex, and medulla. The T cell response, the cell-mediated arm of adaptive immunity that results in direct killing of, e.g., infected or cancerous cells, is coordinated in the paracortex. The B cell response, our humoral or antibody-mediated immunity, is coordinated in the cortex. Lymph fluid travelling from peripheral tissues to the lymph nodes flows around the lymph node and into the inner medulla as well as into the subcapsular sinus, and into the cortex and paracortex. Macrophages that line the subcapsular sinuses break down larger materials into smaller antigens to be released into the cortex or paracortex to stimulate T and B cell responses, and small materials <100 kDa in size enter the cortex and paracortex via the conduit system^1^. The conduit system, also called the reticular network, is made up of fibroblastic reticular cells (FRCs) that ensheath a network of collagen fibers that build a key structural component within the lymph node. FRCs maintain these fibers and produce additional collagen during lymph node expansion and contraction that occur during immune responses^2,3^. Other non-lymphocytes, including fibroblasts, blood and lymphatic endothelial cells, as well as macrophages, also contribute to changes in the lymph node structure during inflammation and as we age.

During inflammatory processes, the lymph node undergoes significant structural changes. Within a few hours of an immune insult, whether a bacterial infection due to a cut or the onset of a flu infection, a massive influx of migrating lymphocytes and antigen presenting cells, including dendritic cells (DCs) coming from the injured tissue, enter the lymph node. At the same time, activated B and T cells and stromal cells (FRCs and lymphatic and blood endothelial cells) begin proliferating^4,5^. During this initial phase, migratory DCs cause FRCs to relax and retract from, or let go of, the collagen fibers, such that the collagen fibers bear most of the tension due to expansion in size, and it is thought that the mesh spacing within the reticular network is increased^7–10^. As the lymph node continues to expand, breaks occur in the collagen fibers and other extracellular matrix (ECM) materials, requiring repair and transfer of some of the tension to the FRCs themselves^11^. Researchers have found permanent changes after the immune response is resolved and the tissue is again at homeostasis because the outer capsule of the lymph node is thicker and stiffer^9^.

Research has shown that the biomechanics, or tension and relaxation that also occur during inflammation in the lymph nodes, can affect cell functions. The FRCs and collagen fibers are hypothesized to provide a structural scaffold for B and T cells that regulates their proliferation and migration, and also to distribute chemokines, cytokines, and growth factors throughout the lymph node^12–15^. T cells respond to their environment by sensing and adapting to the biomechanics, including fluid-like viscous and solid-like elastic stresses. Recently, researchers demonstrated that T cells have higher expression of activation and inhibitory markers when exposed to slow relaxing, more solid-like matrices, while T cells on fast relaxing, more fluid-like matrices had higher expression of memory markers^16^. Biomechanics and viscoelasticity, the combined solid- and fluid-like responses of a tissue, are largely dependent on ECM composition^19^. As we age, we accumulate ECM materials in the lymph nodes (also known as fibrosis), and research has shown that the motility of naïve T cells (circulating T cells prior to exposure to inflammatory stimuli) is decreased in aged compared to young lymph nodes^17^. This is particularly evident near highly ECM-rich fibrotic regions, which are likely to have increased solid-like elastic properties^17^. Aging also appears to modulate naïve T cell numbers, which suggests that fibrosis and biomechanical properties of the lymph nodes could be a contributing factor to age-related changes in our adaptive immune response ^18^. Interestingly, ECM composition and biomechanics of tissues is sex-dimorphic in many organs^20–22^, and these differences may be a contributing factor to sex differences in susceptibility and progression of numerous diseases^20,23^.

Despite the importance of ECM and viscoelasticity in the context of lymphocyte immunity, lymph node biomechanics and viscoelasticity have yet to be examined extensively, including in the context of inflammation. This is in part due to limitations to the methods available to study viscoelastic properties, traditionally via rheology^24^ or atomic force microscopy^25,26^. A rheometer applies a shear or linear stress with a known displacement or speed on the sample and the resulting torque (or stress) the material responds with is recorded. While this is a satisfactory method for homogeneous substances, it is inadequate for samples like the lymph node since the spatial organization vital to its various functions is lost. Additionally, the rheometer usually requires a larger volume than what is available from smaller organs like the lymph nodes. Atomic force microscopy uses a cantilever that touches a sample surface at a constant force. The surface topography causes deflections in the cantilever, which alters the amount of laser light reflected from its surface and captured by a photodetector^25,26^. This readout can be used to infer the sample’s viscoelastic response. While this method preserves the spatial organization of tissues, the analysis is limited to the sample surface and is sacrificial, making it less than ideal for studies aiming to look at both structure and function. Researchers have also used more specific methods, including tension nanoprobes to study lymph node elasticity^27^, tensiometers to study capsule rigidity^28^, 2-photon microscopy to study lymph node fibrosis^17^, and nanoindentors to test lymph node viscoelasticity^9,29^. These methods have been helpful in elucidating lymph node biomechanics, but none of these are non-sacrificial nor provide biomechanical properties beyond the surface.

Using a previously published methodology^30,31^, we combine nanoparticle-based multiple particle tracking (MPT) microrheology with live lymph node slice culture, which allows us to preserve tissue architecture ex vivo^32^ while simultaneously studying lymph node viscoelasticity^30^. MPT uses nanoparticles of sizes close to, but smaller than, the mesh spacing of hydrogel-like tissues and tracks their Brownian diffusive motion within the tissue over time^69–71^. The resulting mean-squared displacement for each particle and the ensemble of all particles is then used to determine biomechanical properties such as elastic and loss moduli, pore size, and microviscosity of the lymph nodes^30,33–36^. Here, we investigate how the biomechanics of the lymph node change during inflammation, particularly in the cortex and paracortex (B and T cell zones). We correlate our microrheology measurements to structural changes in lymph node ECM and assess differences in biomechanical responses to inflammation based on sex. Our study presents, for the first time, an in-depth analysis of the complex biomechanics of distinct lymph node regions during an inflammatory response in the context of tissue remodeling and sex, and lays the foundation for future structure-function relationships between tissue biomechanics and lymphocyte functions.

## RESULTS

### LPS induces peak inflammation in lymph nodes at day 3 and returns to homeostasis by day 14

We inflamed the lymph nodes via intradermal injections of 10 µg lipopolysaccharide (LPS) in female mice to simulate a bacterial challenge. After a single LPS challenge, lymph node mass peaked on day 3 to nearly two-fold compared to untreated mice, started to decrease by day 5 (p<0.1), and returned to masses similar to untreated by day 7 (**Fig. 1A**). We quantified cell numbers in the lymph nodes using flow cytometry (**Supp. Fig. S1C**) and found they first decreased from 4 ± 1 million in untreated to 1.8 ± 0.2 million in day 1 lymph nodes, followed by a significant increase to 4.7 ± 0.7 million on day 3 (**Fig. 1B**). B cells significantly contributed to this increase (**Fig. 1C**), as they increased in number from 0.9 ± 0.2 million cells in untreated to 2.2 ± 0.5 million cells in day 3 lymph nodes (**Fig. 1C**). B cell numbers returned close to values for untreated mice by day 7. While not significantly different, we observed that CD8+ and CD4+ T cell numbers also appeared to contract at day 1 and expand again on day 3 (**Fig. 1D-F**).

**Figure 1.**
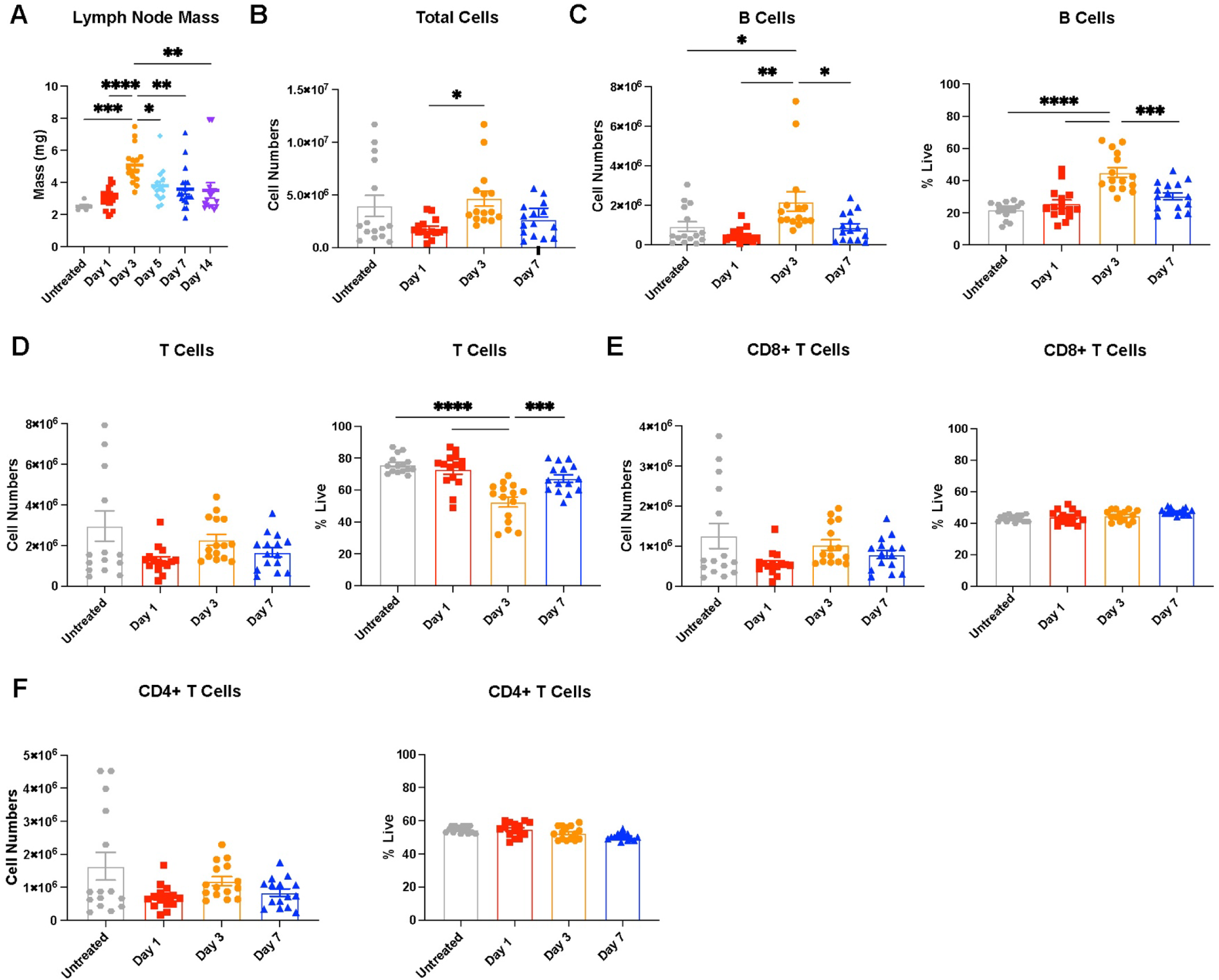
Lymph nodes (LNs) expand and contract over 14 days post single stimulation with lipopolysaccharide (LPS) (A) Mass of inguinal lymph node. (B) Total cell, (C) B cell (CD45+B220+), and (D) T cell (CD45+CD3+) numbers in lymph nodes collected by flow cytometry. (E) CD8+T cell (CD45+CD3+CD8+) and (F) CD4+ T cell (CD45+CD3+CD4+) numbers in lymph nodes collected by flow cytometry 1,3, and 7 days post LPS stimulus. All values are reported as mean ± SEM. Statistical analysis performed by one-way ANOVA followed by Tukey’s post-hoc test **(A-F)**. *p<0.05, **p<0.01, ***p<0.001, ****p<0.0001. N=10-15 female mice per group.

### Peak inflammation increases biomechanical properties of the lymph nodes while reducing pore size

We used multiple particle tracking (MPT) to assess lymph node biomechanical properties, including elastic and loss moduli, pore size, and microviscosity. After performing MPT, the nanoparticles’ mean squared displacement that occurs due to their Brownian motion in the lymph node tissues can be used to extrapolate these various properties, and tissues typically exhibit both elastic and viscous properties. The elastic modulus describes the solid-like properties of a material and refers to how well it returns to its original shape after a force is applied (**Fig. 2A**). The loss modulus describes the liquid-like properties of a material, or the viscous response, that permanently deforms the sample (**Fig. 2A**). We used collagen III and B220 staining to delineate B and T cell zones in the lymph nodes (**Fig. 2B**). We found that 500 nm nanoparticles freely diffuse through the lymph node microenvironment, suggesting that this is the optimal size to probe the ‘mesh’ components, including cells and ECM, as well as fluid components, including interstitial fluid (**Supp. Fig. S1A-B**). We observed a significant increase in lymph node microviscosity during early stages of inflammation, with the overall viscosity reaching a peak value of 3.5 ± 1.2 Pa*s on day 3 (**Fig. 2C)**. Lymph nodes had a significantly higher elastic modulus of 2.2 ± 0.7 Pa on day 3, indicating a stiffer lymph node, compared to all other days (**Fig. 2D, 2E**). Lymph nodes begin to return to elastic modulus values close to healthy by day 7 (0.2 ± 0.06 Pa) (**Fig. 2D-E**). We also observed a significant increase in loss modulus on day 3, 1.9 ± 0.6 Pa, compared to untreated, 0.07 ± 0.02 Pa, which indicates a high resistance to fluid flow at peak inflammation (**Fig. 2D-E**).

**Figure 2.**
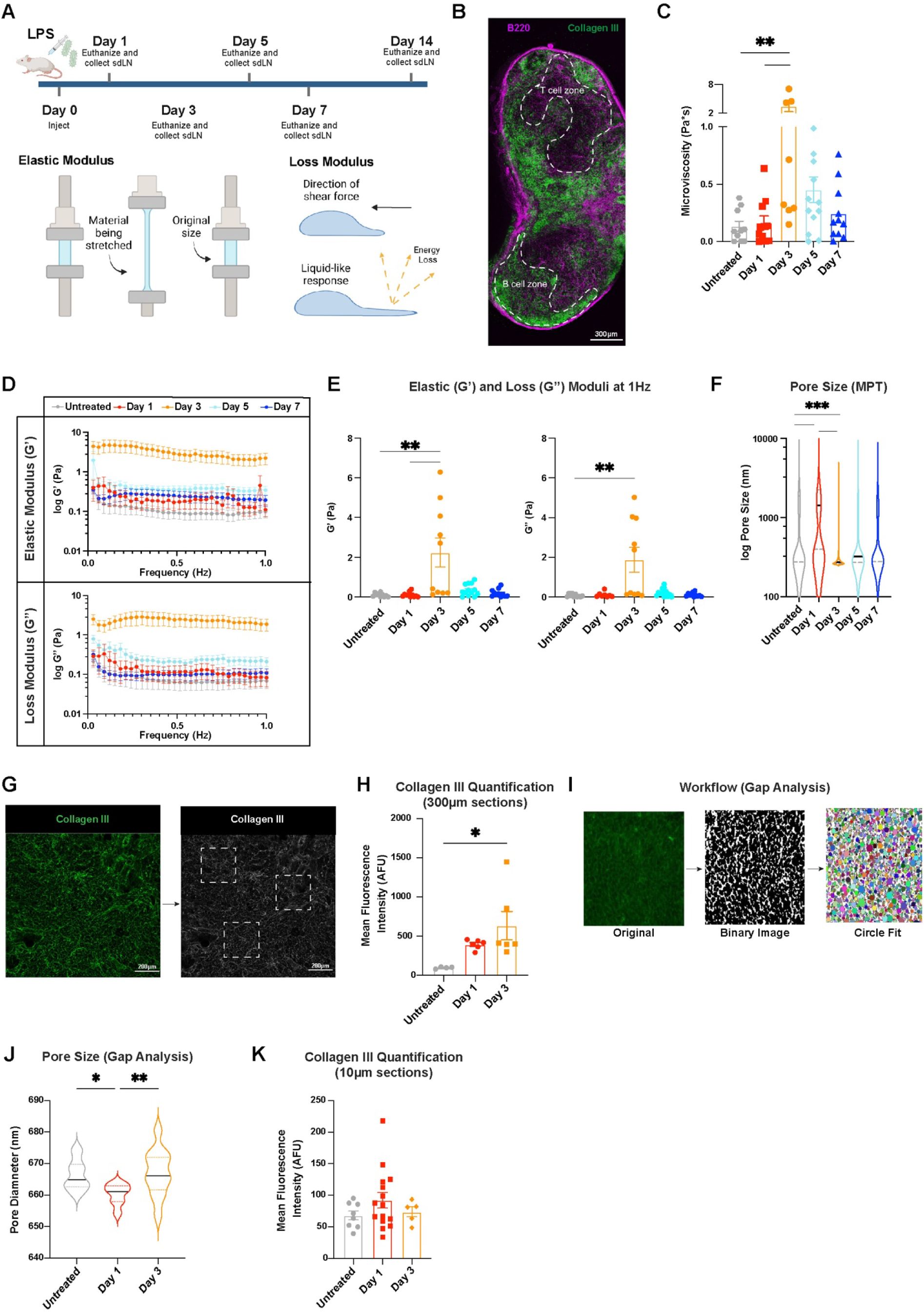
Lymph node biomechanics change in response to LPS. (A) Injection schedule, schematic of elastic modulus, schematic of loss modulus (made with BioRender). (B) Lymph node slice stained for B220 (B cells) and Collagen III (ECM). (C) Microvisccsity of the overall lymph node. (D) Elastic and loss moduli of overall lymph node over 1 Hz and (E) at 1 Hz. (F) Pore sizes (by MPT) in the overall lymph node. (G) Collagen III quantification workflow and (H) Collagen III quantification from confocal images using tissue slices (300µm thick) obtained using a vibratome. (I) Workflow of Gap analysis and (J) pore sizes measured by Gap analysis (PDPN staining). (K) Collagen III quantification using tissue slices obtained via cryosectioning (10µm thick). Median and quartile values shown for pore sizes. Other values are reported as mean ± SEM. Y-axis in **D, F** shown on a logarithmic scale, axes labeled with ‘log’ to enhance readability. Statistical analysis performed with 1-way ANOVA/Kruskal-Wallis tests followed by Tukey’s/Dunn’s multiple-comparisons post-hoc tests **(C,F,H,J,K).** Elastic and loss moduli are compared at 1 Hz as an average across all mice, and statistical analysis was done using Kruskal-Wallis test with Dunn’s post-hoc analysis **(E)**. *p<0.05, **p<0.01, ***p0.001, ***p<0.0001. N = 10-15 female mice.

We found a significant increase in median pore size on day 1 of 1400 nm compared to 800 nm in untreated lymph nodes, indicating tissue relaxation at the start of the inflammatory response (**Fig. 2F**). Pore size then decreased significantly to 270 nm on day 3 (**Fig. 2F**) and began to increase again thereafter. Our data indicate that changes in lymph node elastic and loss moduli are closely related to changes in cell proliferation during acute inflammation.

To understand underlying causes for changes in pore size, we semi-quantitatively assessed the key ECM component in the lymph node, collagen III, via fluorescence microscopy (**Fig. 2G**). We used either 10µm (obtained via cryostat) or 300µm (obtained via vibratome) thick lymph node sections. In the thicker vibratome slices, we found a significant increase in collagen III mean fluorescent intensity (MFI) from 98 ± 6 AFU in untreated to 640 ± 180 AFU on day 3 (**Fig. 2H**). This data suggests that increases in collagen density may contribute to the decreased pore size observed on day 3 (**Fig. 2F**). In addition, using thin, 10µm lymph node slices stained for podoplanin, a marker of FRCs, in conjunction with a MATLAB script, Gap analysis^7^, we calculated pore sizes within the reticular network. Gap analysis fits circles of the largest possible diameter into binarized images of stained tissue to identify and measure non-overlapping gap sizes (**Fig. 2I**). The pore sizes obtained using this method showed different trends to those obtained from MPT data: the median pore size reduced slightly from 670 nm in untreated to 660 nm on day 1 and increased again on day 3 to 670 nm (**Fig 2J**). These trends were likely different since, unlike MPT, Gap analysis does not take into consideration presence of the effects due to cells or other non-collagen materials during the analysis of pore size. We also found no differences in collagen MFI in thin sections (**Fig 2K**). These contrasts highlight that ‘pore size’ or ‘mesh spacing’ is not a uniform metric; the characterization method used can substantially influence both the measured values and their interpretation in hydrogel-like tissues.

### B and T cell zones exhibit similar biomechanical responses during acute inflammation

B and T cell zones, or cortex and paracortex, in the lymph nodes serve unique functions that lead to B and T cell activation during an inflammatory response. Given the distinct functional roles of B and T cell zones, we next assessed whether their biomechanical properties differed. Surprisingly, the microrheological profiles of the two compartments were similar. Mean microviscosity values were comparable, ranging from 0.055 – 2.3 Pa*s in the B cell zone to 0.16 – 3.8 Pa*s in the T cell zone (**Fig. 3A**), with both reaching a peak at day 3. Notably, despite these similarities, individual measurements varied considerably between mice (**Supp. Fig. S2A**), highlighting substantial inter-animal heterogeneity. Similarly, we found that trends in elastic and loss moduli were comparable between both zones, with both reaching a maximum value at day 3 **(Fig 3B-C).** However, direct comparisons at each time point (**Supp. Fig. S2B**) showed no significant differences between the two compartments. These findings, together with the microviscosity data, indicate that B and T cell zones exhibit similar biomechanical behavior during acute inflammation.

**Figure 3.**
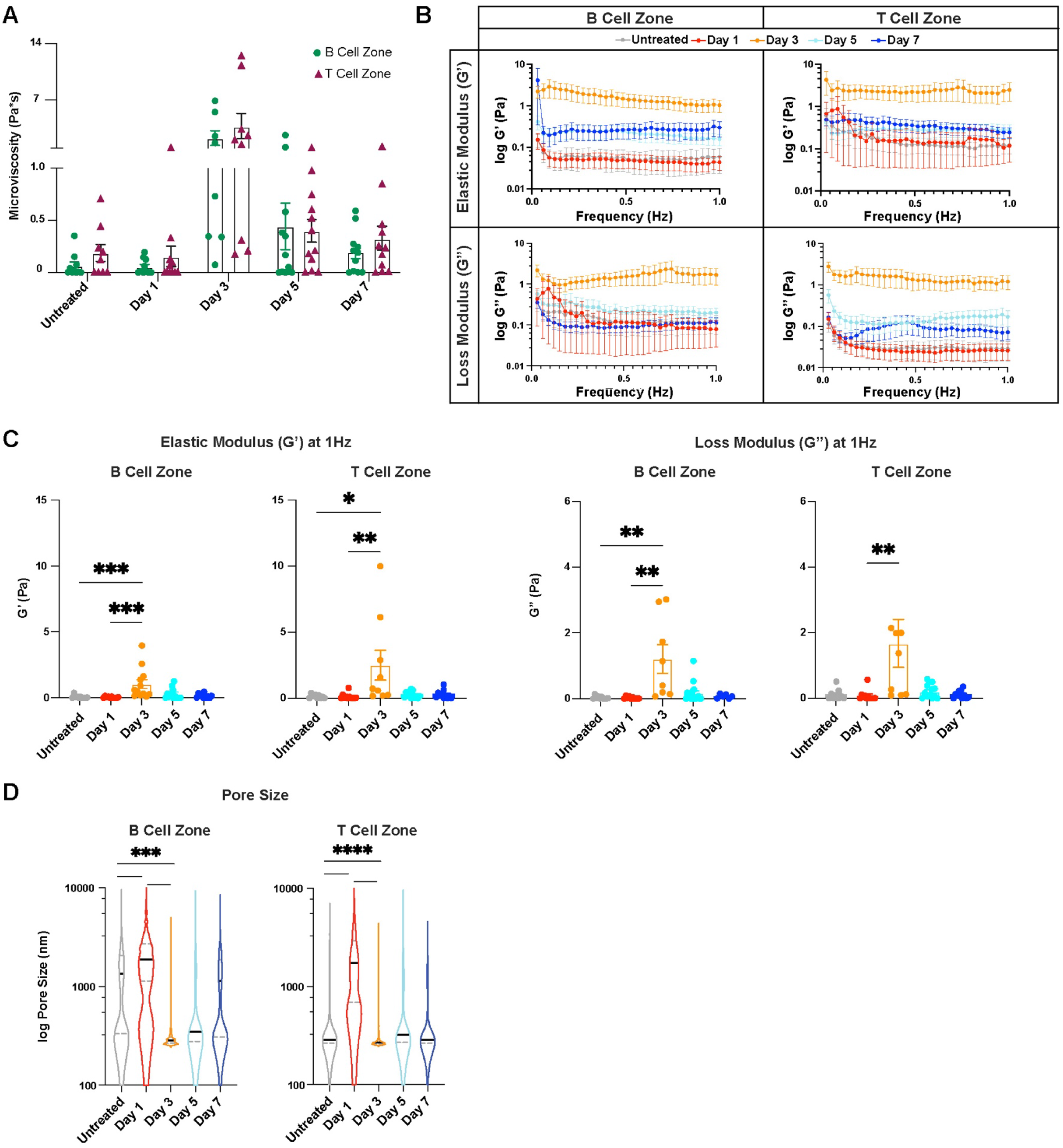
B and T cell zone biomechanics change in a similar fashion during immune response to LPS. (A) Microviscosity of the B and T cell zones in untreated lymph nodes and lymph nodes days 1,3,5, and 7 after LPS treatment. (B) Elastic and loss moduli of B and T cell zones over 1 Hz and (C) at 1 Hz. (D) Pore sizes (by MPT) of B and T cell zones during the course of inflammation. Median and quartile values shown for pore sizes. Other values are reported as mean ± SEM. Y-axis in **B, D** shown on a logarithmic scale, axes labeled with ‘log’ to enhance readability. Statistical analysis of zone-wise comparison of microviscosity was performed using a Mann-Whitney test on each day **(A)**, elastic and loss moduli are compared at 1Hz as an average across all mice, and statistical analysis is was performed with a Kruskal-Wallis test with Dunn’s post-hoc analysis **(C)**. Statistical analysis for pore size was done using a Kruskal-Wallis test followed by Dunn’s post-hoc test **(D)**. *p<0.05, **p<0.01, ***p<0.001, ****p<0.0001. N = 10-15 female mice.

In both the B and T cell zones, there was an initial increase in pore size. In the B cell zone, median pore size increased from 1300 nm in untreated to 1900 nm on day 1 and then decreased to 280 nm on day 3, followed by an increase to 350 nm on day 5 and 1100 nm on day 7 (**Fig. 3D**). Similarly, the T cell zone median pore size increased from 290 nm in untreated to 1700 nm on day 1 and then decreased to 270 nm on day 3 and remained relatively stable at 320 nm on day 5 and 290 nm on day 7, similar to the pore size of untreated lymph nodes (**Fig. 3D**).

### Lymph node biomechanics change permanently after resolution of acute inflammation

Two weeks after inflammation is induced, the overall cell numbers in the lymph node have contracted to 2 ± 0.6 million cells, similar to levels in untreated lymph nodes (4 ± 1 million cells, (**Fig. 4A**). Similarly, B cell and T cell numbers (**Fig. 4B**), including CD4+ and CD8+ T cell subsets (**Fig. 4C**), return to levels similar to untreated. These data suggest that at day 14, lymph node cell composition has returned to states similar to untreated, and resolution has occurred. However, we have also found that biomechanical properties of the lymph nodes do not fully return to baseline. Lymph node microviscosity increased 3.5-fold on day 14 compared to untreated (**Fig. 4D**). Though not significantly different, B cell zone microviscosity appeared to be lower than T cell zone microviscosity for both untreated and day 14 (**Fig. 4D**). We found a higher elastic modulus, 0.3 ± 0.09 Pa, and loss modulus, 0.2 ± 0.07 Pa, on day 14 compared to 0.1 ± 0.03 Pa and 0.07 ± 0.02 Pa for untreated, respectively, (**Fig. 4E-F**), indicating an overall stiffer tissue that had higher resistance to flow. These data suggest that lymph nodes on day 14 are permanently restructured after a single immune challenge. B cell zones appeared to drive much of these changes, as a greater increase on day 14 was observed for both loss and elastic moduli compared to T cell zones (**Fig. 4E**). We also found a significant decrease in median pore size on day 14 (350 nm) compared to untreated (800 nm) (**Fig. 4G**). Additionally, we found an increase in collagen III on day 14 in thick sections compared to untreated **(Fig. 4H).** When assessing pore size based on staining only for the FRC network (podoplanin) using Gap analysis, we also found a slight reduction in pore size at day 14 compared to untreated, and thin sections revealed no differences in collagen III **(Fig. 4I-J).** Differences in the values of pore size and collagen quantifications for these two methods are likely due to considering a larger (300 µm MPT/thick sections) or smaller (10 µm Gap analysis/cryosections) section of the lymph nodes.

**Figure 4.**
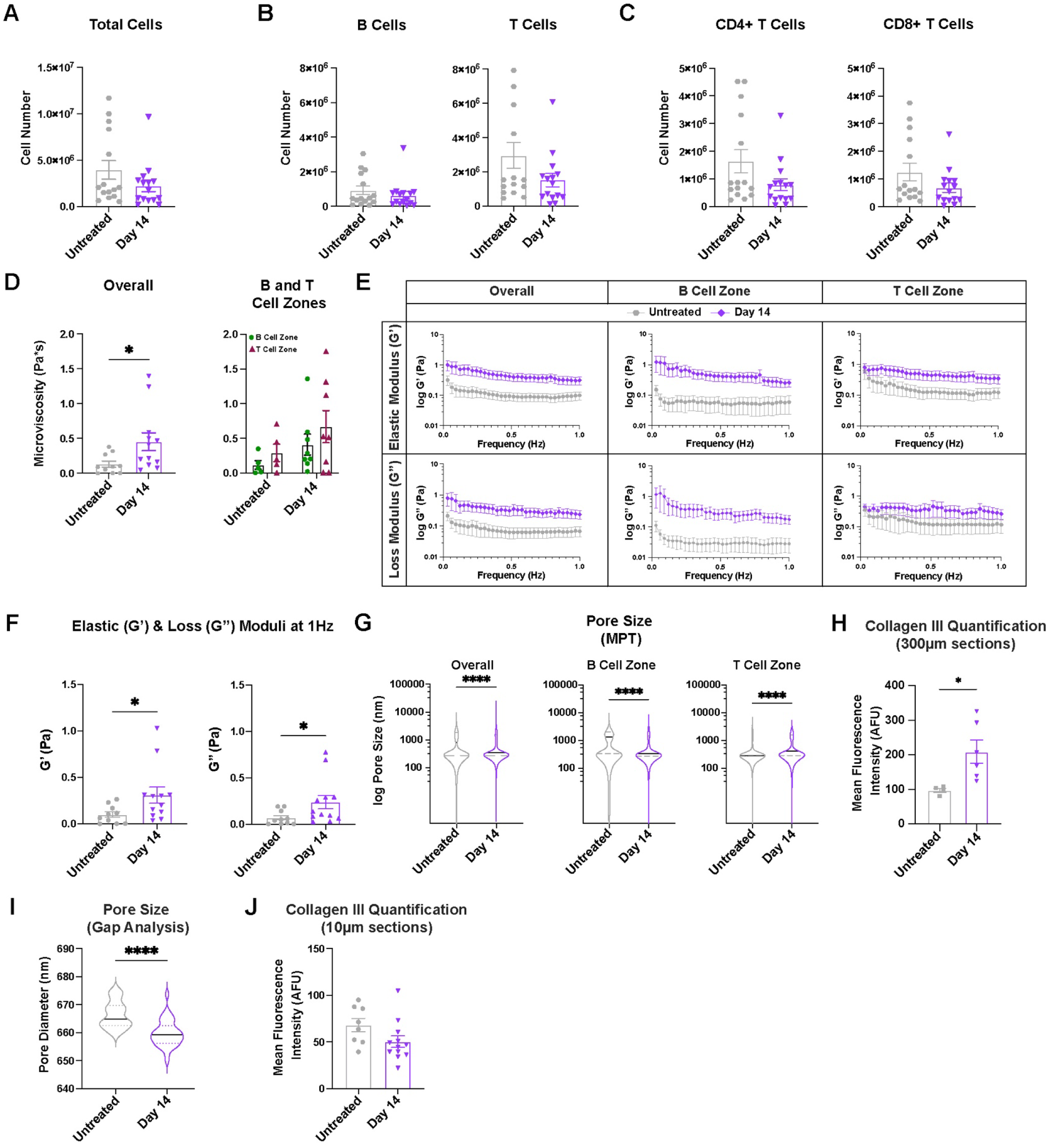
Lymph node biomechanics show permanent changes at resolution, two weeks after acute inflammatory challenge with LPS. (A) Total cell, (B) B cell (CD45+B220+),T cell (CD45+CD3+), and (C) CD8+ T cell (CD45+CD3+CD8+) and CD4+ T cell (CD45+CD3+CD4+) numbers in lymph nodes from untreated mice and mice 14 days post LPS stimulation analyzed by flow cytometry. (D) Microviscosity in the overall lymph node and in the B and T cell zones from untreated mice and mice 14 days post LPS stimulation. Elastic and loss moduli over (E) 1 Hz and (F) at 1 Hz in the overall lymph node and in the B and T cell zones from untreated mice and mice 14 days post LPS stimulation. (G) Pore sizes (by MPT) in the overall lymph node and in the B and T cell zones. (H) Collagen III quantification from confocal images using tissue slices obtained via vibratome (300µm thick) (I) Pore sizes measured by Gap analysis (PDPN staining). (J) Collagen III quantification using tissue slices obtained from a cryostat (10µm thick). Median and quartile values shown in pore sizes. Other values are reported as mean ± SEM. Y-axis in **E, G** shown on a logarithmic scale, axes labeled with ‘log’ to enhance readability. Statistical analysis performed by a t-test/Mann-Whitney test **(A-D, G, H, I).** Statistical analysis of zone-wise comparison of microviscosity was performed with a Mann-Whitney test at each timepoint **(D)**. Elastic and loss moduli are compared at 1 Hz as an average across all mice, and statistical analysis is done via Mann-Whitney test **(E,F)**. *p<0.05, **p<0.01, ***p<0.001, ****p<0.0001 N = 10-15 female mice.

### Changes in lymph node biomechanics during inflammation are sexually dimorphic

To determine whether biological sex influences lymph node biomechanics during an acute inflammatory response, we compared the biophysical properties of the lymph nodes between male and female mice at days 3 and 14 post injection, in addition to untreated, to assess differences at peak inflammation and resolution, respectively. We found that female lymph nodes exhibit slightly higher elastic moduli already in untreated states, with 0.11 ± 0.03 Pa in female mice and 0.05 ± 0.004 Pa in male mice (**Fig. 5A-B**), though not significant. At day 3, lymph nodes had significantly higher elastic moduli in female compared to male mice, with 2.2 ± 0.7 Pa vs. 0.01 ± 0.003 Pa, respectively. On day 14, lymph nodes from female mice continued to exhibit higher elastic moduli than those from male mice with 0.3 ± 0.08 Pa vs. 0.05 ± 0.01 Pa (**Fig. 5A-B**), respectively. We found similar results for loss moduli, where the moduli in lymph nodes from female mice were higher compared to those from male mice for both day 3, with 1.9 ± 0.6 Pa (female) vs. 0.005 ± 0.001 Pa (male), and day 14, with 0.23 ± 0.07 Pa (female) vs. 0.02 ± 0.006 Pa (male) (**Fig. 5A-B**).

**Figure 5.**
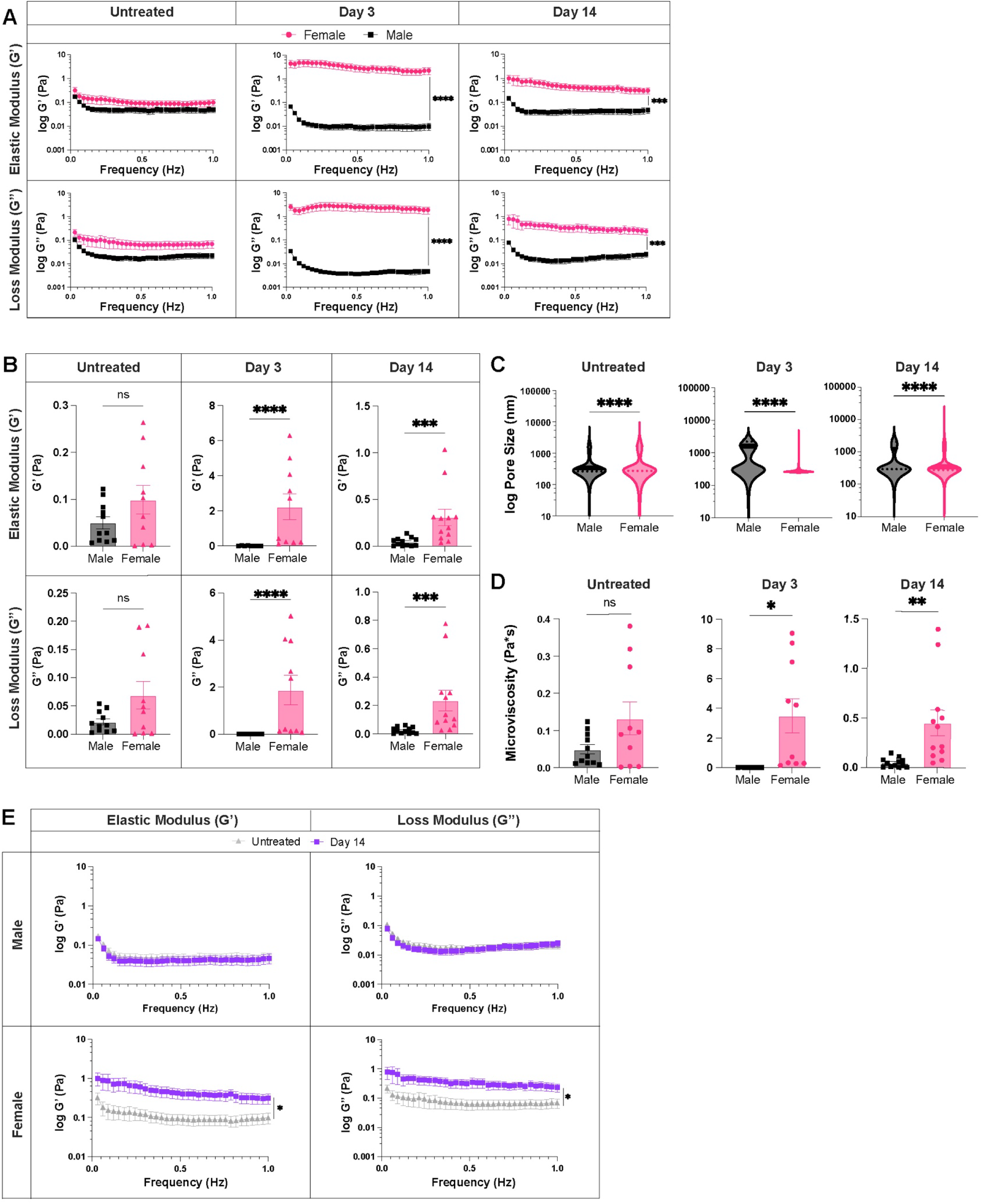
Lymph node biophysical properties are sexually dimorphic. Elastic and loss moduli for male and female lymph nodes (A) over 1 Hz and (B) at 1 Hz. (C) Overall pore size (by MPT), and (D) microviscosity of healthy and acutely inflamed male and female lymph nodes. (E) Elastic and loss moduli of male and female lymph nodes for untreated and day 14 lymph nodes over 1 Hz. Median and quartile values shown in pore sizes. Other values are mean ± SEM. Y-axis in **A, C, E** shown on a logarithmic scale, axis labelled with ‘log’ to enhance readability. Statistical analysis by Mann-Whitney test (**B,C**) or Welch’s t-test (**D**). Elastic and loss moduli are compared at 1 Hz as an average across all mice, and statistical analysis is done by t-test/Wilcoxon rank-sum test(**A,B,E**). Significance values are denoted on **A,E** for representation.*p<0.05, **p<0.01, ***p<0.001, ****p<0.0001, ns p≥0.05. N=8-12 male or female mice.

We also found that while elastic moduli and loss moduli reach a peak at day 3 in female mice, they reach a minimum in male mice (**Fig. 5A**). Additionally, we found lower median pore sizes (270 nm vs 1600 nm) (**Fig. 5C**) and higher mean viscosity (3.5 ± 1.1 Pa*s vs. 0.009 ± 0.002 Pa*s) (**Fig. 5D**) on day 3 in lymph nodes from female vs. male mice. Similar trends were observed within the B and T cell zones in lymph nodes of male and female mice at all time points (**Supp. Fig. S3A-C, S4A-B).** Additionally, like female mice, there were no biomechanical differences observed between the B and T cell zones in male mice. When assessing lymph node biomechanics at resolution of inflammation, we found a return to baseline values for both elastic moduli (0.05 ± 0.01 Pa vs. 0.05 ± 0.01 Pa) and loss moduli (0.02 ± 0.005 Pa vs. 0.02 ± 0.006 Pa) in male mice **(Fig 5E)**. However, for female mice, both elastic (0.3 ± 0.08 Pa at day 14 vs. 0.1 ± 0.03 Pa in untreated) and loss moduli (0.23 ± 0.07 Pa at day 14 vs. 0.07 ± 0.02 Pa in untreated) were increased at day 14 (**Fig. 5E**). Independent B and T cell zone restructuring was observed only in lymph nodes from female (**Fig. 4E-F)** mice (**Supp. Fig. S4C).** These data suggest that biological sex influences lymph node restructuring during and after resolution of acute inflammation.

### Chronic inflammation does not modulate lymph node biomechanics

Given our findings that acute inflammation induces changes in lymph node biomechanics, we next sought to understand if these changes are enhanced during chronic inflammation. To induce chronic inflammation, 10 µg LPS was intradermally injected weekly for 5 weeks and mice euthanized days 3 and 14 after the final injection were chosen to assess lymph node biomechanics (**Fig. 6A**). Chronically inflamed lymph nodes trended toward more cells on days 3 and 14 (3 ± 0.8 million cells and 2 ± 0.7 million cells, respectively), compared to untreated (0.5 ± 0.09 million cells) (**Fig. 6B**). Compared to untreated (0.1 ± 0.01 million), B cell numbers were increased on day 3 (1 ± 0.4 million) and day 14 (0.6 ± 0.2 million) (**Fig. 6C**). Similarly, there was a trend toward total T cell numbers on day 3 (1.6 ± 0.3 million) and day 14 (1.2 ± 0.4 million) compared to untreated (0.4 ± 0.1 million) (**Fig. 6C**). CD4+ and CD8+ T cell numbers also trended higher on day 3 (0.9 ± 0.2 million and 0.6 ± 0.01 million, respectively) compared to day 14 (0.7 ± 0.2 million and 0.4 ± 0.2 million) (**Fig. 6D**). While the total numbers of total cells, B cells, and T cells were only trending toward significant (p values for all were p<0.15), the percent live of each of these was significantly higher at day 3 compared to untreated. By day 14 after the final injection, all cell numbers decreased again and were not significantly different to untreated.

**Figure 6.**
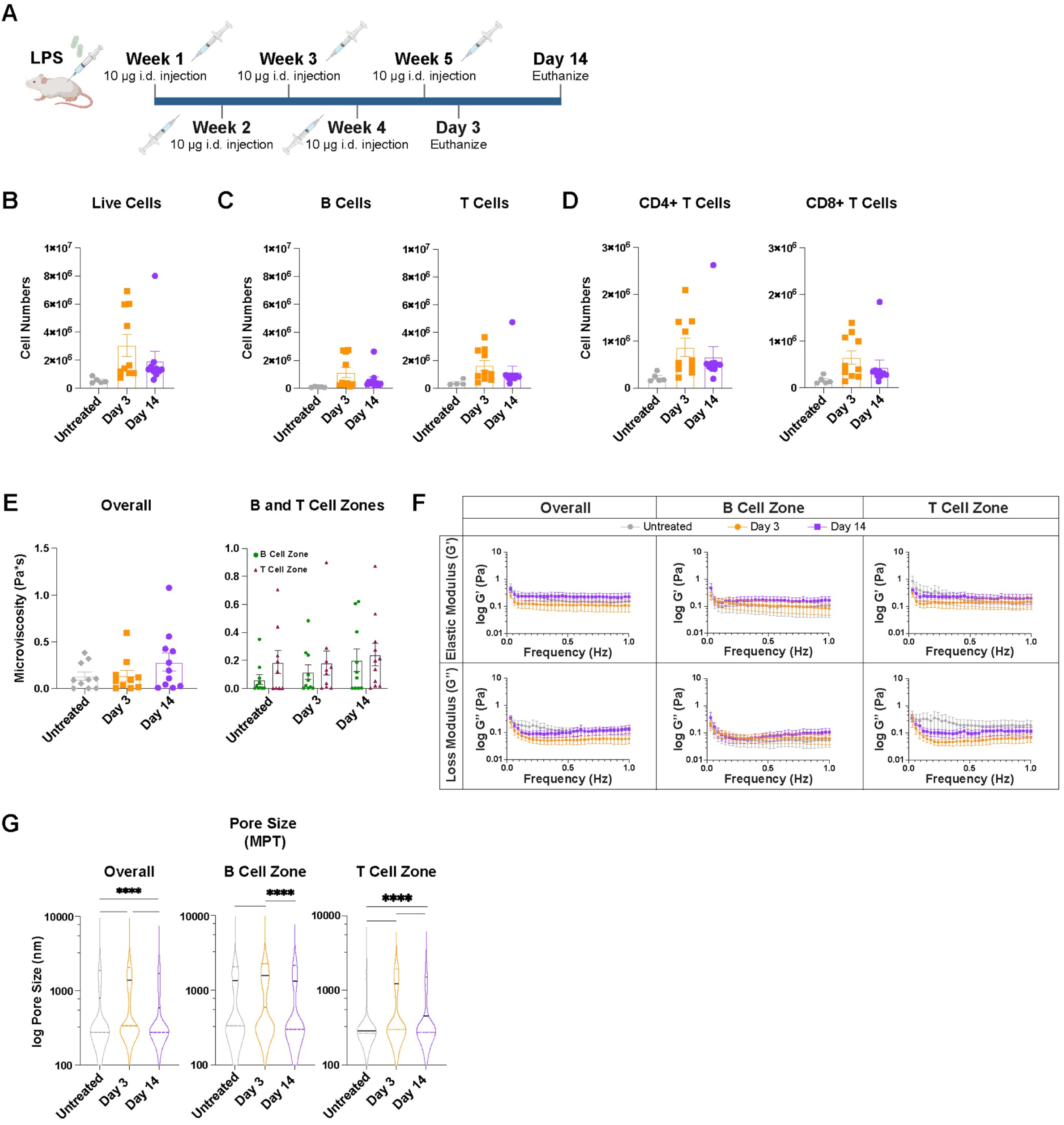
Chronic inflammation does not modulate lymph node biomechanics. (A) Injection timeline for chronic inflammation studies (B)Total cell, (C) B cell (CD45^+^B220^+^) and T cell (CD45^+^CD3^+^), and (D) CD8+ T cell (CD45^+^CD3^+^CD8^+^) and CD4+ T cell (CD45^+^CD3^+^CD4^+^) numbers from untreated and chronically inflamed lymph nodes analyzed by flow cytometry. (E) Micro viscosity (F) elastic and loss modulus over 1 Hz, and (G) pore sizes (by MPT) in the overall lymph node and in the B and T cell zones of untreated chronically inflamed lymph nodes. Median and quartile values shown in pore sizes. Other values are reported as mean ± SEM. Y-axis in **F,G** shown on a logarithmic scale, axis labelled with ‘log’ to enhance readability. Statistical analysis by 1-way ANOVA/Kruskal-Wallis test followed by Tukey’s/Dunn’s multiple-comparisons post-hoc test **(B-D, E, G).** Statistical analysis of zone-wise comparison of microviscosity was done by a Mann-Whitney test on each day **(E)**. Elastic and loss moduli are compared at 1Hz as an average across all mice, and statistical analysis is done by Kruskal-Wallis Test **(F)**. *p<0.05, **p<0.01, ***p<0.001, ****p<0.0001. N = 10-15 female mice.

Biomechanical properties of chronically inflamed lymph nodes appeared to be similar 3 days and 14 days after final injection and remained similar to untreated. Chronically inflamed lymph nodes had microviscosities of 0.1 ± 0.04 Pa*s for untreated, 0.1 ± 0.06 on day 3, and 0.3 ± 0.1 Pa*s on day 14 (**Fig. 6E**). T and B cell zones also had similar microviscosities on day 3 (0.2 ± 0.08 and 0.1 ± 0.05 Pa*s, respectively) and day 14 (both 0.2 ± 0.08 Pa*s) (**Fig. 6E**). Elastic moduli in lymph nodes were also similar on day 3 (0.1 ± 0.05 Pa*s), day 14 (0.2 ± 0.08 Pa*s), and for untreated (0.09 ± 0.03 Pa) (**Fig. 6F**). Elastic moduli were slightly lower on day 3 compared to day 14 in both B (0.08 ± 0.04 Pa and 0.2 ± 0.06 Pa, day 3 and 14, respectively) and T cell zones (0.15 ± 0.07 Pa and 0.2 ± 0.08 Pa, day 3 and 14, respectively) (**Fig. 6F**), though this was not significant. Loss moduli were similar on day 3 (0.06 ± 0.02 Pa*s) and day 14 (0.1 ± 0.05 Pa*s) compared to untreated (0.07 ± 0.02 Pa) (**Fig. 6F**). Loss moduli were also similar at days 3 and 14 in B (0.06 ± 0.03 Pa and 0.1 ± 0.04 Pa, day 3 and 14, respectively) and T cell zones (0.07 ± 0.02 Pa and 0.1 ± 0.04 Pa, day 3 and 14, respectively) (**Fig. 6F**). Median pore size within the lymph node significantly increased from 800 nm in untreated to 1400 nm on day 3 (**Fig. 6G)**; this was true for both B (1300 nm in untreated to 1600 nm at day 3) and T (300 nm in untreated to 1200 nm at day 3) cell zones (**Fig. 6G)**. These patterns remained true one month (“long-term resolution”) after the final LPS challenge. Median pore sizes within the lymph node were significantly increased from untreated (800 nm) to long-term resolution (1200 nm) (**Supp. Fig. S5A**), as well as in the B (1300 nm untreated to 1400 nm long-term) and T (300 nm untreated to 1100 nm long-term) cell zones (**Supp. Fig. S5A**). Elastic and loss moduli remained similar to untreated at long-term resolution both overall and in the B and T cell zones **(Supp. Fig. S5C)**.

## DISCUSSION

Elucidating the role of the influence of biomechanical stimuli on immune cell behavior and function has been an area of active interest^37–39^. In this study, we examined the biomechanical properties of the lymph node, a central pillar involved in the coordination of immune response. Using multiple particle tracking (MPT) microrheology^30,31,36^, a minimally invasive technique, we provide a comprehensive analysis of lymph node biomechanics (elastic/loss moduli, pore size, microviscosity) during homeostasis and over the course of acute and chronic inflammation, including at resolution. We found that biomechanics of B and T cell zones change in a similar fashion during acute inflammation, and that changes in biomechanical properties are sex dependent. Our work suggests that restructuring may in part depend on changes in extracellular matrix (ECM) composition, and that chronic inflammation does not cause the same differences in biomechanical response as acute inflammation.

Lipopolysaccharide (LPS) initiates a TLR-4 response, which resulted in a lymph node expansion-contraction cycle that lasted approximately 14 days. Lymph nodes increase in size as soon as 1 day post initial stimulus and reach an inflammation peak at day 3 after LPS stimulation. We observed that an initial increase in the pore size at day 1 compared to untreated counterparts matches prior observations that the inter-network spacing of the reticular network is increased during the initial phases of lymph node expansion^7,8,10,11,40^. However, we did not detect a reduction in elastic and loss moduli at day 1, unlike previous reports^6,8,10^ that describe a ‘priming phase’, in which network tension decreases, or ‘relaxes’, during early phases of lymph node expansion. This ‘relaxation’ reflects a shift in load bearing from the conduits to cells in the fibroblastic reticular cell (FRC) network, features which our microrheological approach may not be able to directly resolve. In addition, immune response kinetics to LPS alone may likely differ from those of antigen-adjuvant models used prior, with earlier stages dominated by innate activation, leading to the altered mechanical trajectories we observed. By peak inflammation (3 days post injection), lymph nodes are nearly doubled in size, on account of increased cellularity. Previous studies^4,5,7,9^ have shown that lymphocyte influx plays a key role in lymph node swelling mechanics and that perturbing the entry of lymphocytes leads to both a lowered viscosity and lymph node volume^9^. Lymph node swelling is also associated with stretching of the reticular network and changes to FRC size^8,9,40^. We note the increased cellularity at day 3 corresponds to a decrease in pore size and the higher viscosity echoes trends reported earlier. A consequence of increased cellularity could be a decrease in interstitial fluid density within the lymph node, which coupled with reduced pore sizes, may result in altered cell migration patterns within the lymph node. Smaller pores reduce the ease of nuclear movement for migrating cells^42^, which can have adverse effects on their survival^43,44^ as they navigate through the complex lymph node microenvironment. Interestingly, we observed a high degree of heterogeneity in the rheological data obtained at day 3, likely reflecting both biological and structural variability at peak inflammation. Differences in cell packing due to increased cellularity, variations in FRC network tension, and individual response trajectories across mice^8^ may contribute to the observed rheological differences, despite synchronized sampling. Further, active cellular processes such as cell migration, enzymatic remodeling, or fibrosis can also cause microscale changes in the tissue structure. These changes are spatially and temporally heterogeneous during immune response and can lead to additional variations in the measured microviscosity and moduli. Consequently, while MPT is a powerful tool, future investigations integrating it with other high-throughput biophysical techniques or imaging approaches will help provide a more comprehensive understanding of complex tissue biomechanics.

Quantifying the morphology—particularly the pore size or mesh spacing—of hydrogels and hydrogel-like tissues has been a long-standing interest in tissue engineering, drug delivery, and the study of cell migration^62–66^. The pore size distribution significantly impacts the mechanical properties, such as elastic and loss moduli, of hydrogels and tissues. Therefore, precise determination of pore sizes has broad implications for a wide range of biomedical applications. The term ‘pore size’ is often used as a blanket descriptor, yet its definition and interpretation rely heavily on the measurement method. In our study, we used two approaches, Gap analysis and MPT, to estimate the pore size of lymph node tissue. Gap analysis provides a direct, spatial estimate of pore size, measuring the physical space between elements of the podoplanin-stained reticular network. In contrast, MPT yields an indirect, dynamic, and temporally informed metric based on individual particle movement within the complete tissue instead of a single stained component. This allows it to integrate the influence of cells, ECM, and all tissue components, not just the FRC network, on pore size. Each method has its own limitations: Gap analysis is restricted by a limited field of view and is susceptible to artifacts introduced during sample processing and image stitching. MPT, as a 2D approximation, cannot resolve errors due to 3D particle motion, and is sensitive to noise that may skew particle mobility measurements. The differences in our results highlight that studies of pore size in complex tissues, such as the lymph node, are not always directly comparable, and the characterization method should therefore be explicitly considered when interpreting or generalizing ‘mesh spacing’ in tissues. We also observed a negative correlation between pore size and collagen III deposition, suggesting increased ECM deposition may be responsible for changes in tissue morphology, and consequently tissue biomechanics. Exploring the relative contributions of the ECM on lymph node biomechanics is a key area for subsequent investigation.

Our work also revealed that the B and T cell zones of the lymph node exhibit similar restructuring patterns. The microviscosity and elastic and loss moduli exhibit no differences across the zones during acute inflammation. Interestingly, however, the B and T cell zones exhibit different trends in pore size, and the ratio of B to T cell zone pore sizes shows an increasing trend over the course of an immune challenge, suggesting a higher degree of reorganization in the B cell zones. These data reflect similar patterns observed in other studies that highlight changes in the relative volume fractions of the B cell zone and T cell zone in the lymph node following antigen stimulation - there was a notable increase in the size of the B cell zone with a concurrent decrease in the T cell zone. Differences in restructuring may be due to the maturation of follicles to germinal centers at different time points. B cells undergo somatic hypermutation over time and these cellular changes may affect restructuring. The prevailing view is that the overall changes in lymph node structure were largely due to the reticular network in the T cell zone^9,13,15,45^, but these findings indicate that the B cell zone may play a larger role than previously thought.

The prevalence of sex bias in immune cell counts and immune response between males and females is well documented^46–48^. Here, we demonstrated for the first time that biological sex also influences the mechanical properties and restructuring of lymph node tissue during acute inflammatory response. Lymph nodes in female mice exhibit higher elastic and loss moduli than lymph nodes from male mice over the course of immune response. A higher elastic modulus indicates tissues are more resistant to deformation (stiffer), whereas a higher loss modulus indicates tissues dissipate more energy under deformation, indicating a higher degree of internal friction. From a structural or biochemical perspective, higher elastic and loss moduli could reflect alteration of ECM structure in a sex-specific fashion due to differences in matrix protein concentrations, alignment, or densities^72^. Interestingly, 3 days post inflammatory stimulus, we observe a peak in elastic/loss moduli in female lymph nodes but a minima in male lymph nodes. This reversal of trends could be indicative of males and females exhibiting differing immune response rates to LPS, which may be influenced by sex hormones. Indeed, several clinical studies and in vitro models have demonstrated higher proinflammatory cytokine release^49–52^ in males compared to females. Conversely, there is evidence to suggest that the influence of male sex hormones, androgens, may have an inhibitory effect^53,54^, leading to sex-based discrepancies and a diminished immune response in males. One study suggests that the TLR4 receptor, responsible for detection of LPS, may be regulated by testosterone. They found TLR4 expression on macrophages was reduced in the presence of testosterone in vitro, and surgical castration led to an increase in TLR4 expression and elevated inflammatory response in male mice^55^. Another study^56^ demonstrated that androgens down regulated the number of skin-resident dendritic cells and lowered their migratory capacity to lymph nodes in male mice. While the effect of androgens may provide an explanation for our observations during lymph node expansion, additional research is needed to accurately assess the effect of sex hormones on lymph node biomechanics during the resolution phase of acute inflammation.

Disease states such as chronic inflammation, and even aging, have been shown to alter lymph node structure as well as impair immune function^17,57^. In aging, there is an increase in collagen deposition and a blurring of boundaries between B and T cell zones in the lymph nodes^18,58,59^. This suggests that aging may result in stiffer lymph node tissue environments, and research has shown that viscoelasticity can modulate T cell functions, including expression of activation vs. memory markers^16^. Furthermore, aging appears to reduce T cell migration speed and linearity, which is also exacerbated in regions close to the extracellular matrix, suggesting that the biophysical environment surrounding T cells can affect their functions^17^. One hypothesis in aging is that the repeated inflammatory insults we experience throughout our lives cause permanent lymph node remodeling. We sought to mimic this by inducing chronic inflammation, but our data suggest that at least after one month of inflammation, there are minimal biophysical changes in the lymph nodes. However, our study may be limited by the length and type of inflammatory cues we used ^67,68^. Remodeling of tissues in chronic inflammation may be slower than the time scales we observed; thus, future research investigating how different types of inflammation and duration of chronic inflammatory stimuli impact lymph node biophysical properties would provide valuable insights and enhance the generalizability of our findings.

In summary, the role of biomechanics is heavily understudied in the context of immune responses, particularly in the lymph nodes. This study investigated for the first time how the biomechanics of the lymph node are altered during acute and one-month long inflammation, and highlights the vital role of both the reticular network and lymph node architecture in modulating lymph node biophysical properties over the course of an acute inflammatory response. It also sheds light on the role of biological sex on lymph node biomechanics, providing an additional complexity to known sex differences in immune responses. Our work provides new insights into structure-function relationships in the lymph node and underscores the growing importance of tissue mechanics in understanding immune response.

## MATERIALS AND METHODS

### Animal work

Female and male C57BL/6 mice, 6-10 weeks old were intradermally injected with 10µg of LPS (Invivogen) to stimulate the right inguinal skin draining lymph node. Mice were euthanized by CO_2_ inhalation 1, 3, 5, 7, or 14 days after injection. Lymph nodes were collected and any fat surrounding the tissue was removed. Tissues were then placed in cold 1x phosphate buffered saline (PBS) with 2% fetal bovine serum (FBS, ThermoFisher). All animal work was approved by the Institutional Animal Care and Use Committee.

### Flow cytometry

After intradermal injection with LPS, inguinal lymph nodes were harvested for flow cytometry analysis. A single cell suspension was collected by pushing lymph nodes through a 70 μm cell strainer and then resuspending in RPMI (ThermoFisher) supplemented with 10% FBS (ThermoFisher), 1× l-glutamine (Corning), 50 U mL^−1^Pen/Strep (Sigma), 50 μM beta-mercaptoethanol (ThermoFisher), 1 mM sodium pyruvate (ThermoFisher), 1× nonessential amino acids (Fisher Scientific), and 20 mM HEPES (GE Healthcare). Single cell suspensions were stained for viability (live/dead), B cells (B220+), total T cells (CD3+), helper T cells (CD4+), and cytotoxic T cells (CD8+). Stained cell suspensions were analyzed using flow cytometry with a BD FACS Celesta and FlowJo Version 10.

### Tissue slicing

Slicing of lymph nodes was done as previously described^30,32^. Briefly, tissues were embedded in a 6% w/v low melting agarose (Thomas Scientific) and left to solidify. Once the gel was solidified, tissues were punched out using a 10 mm biopsy punch (Robbins Instruments). Gel pieces were placed on the sample tray of a Leica VT1000s vibratome (speed of 3.9 mm/s, frequency of 0.3 Hz, and amplitude of 0.6 mm) and sliced 300 µm thick. Slices were collected using a brush and placed into RPMI (ThermoFisher) supplemented with 10% FBS (ThermoFisher), 1× l-glutamine (Corning), 50 U mL^−1^Pen/Strep (Sigma), 50 μM beta-mercaptoethanol (ThermoFisher), 1 mM sodium pyruvate (ThermoFisher), 1× nonessential amino acids (Fisher Scientific), and 20 mM HEPES (GE Healthcare). Lymph node slices were incubated at 37 °C with 5% CO_2_ in media for at least 1 h prior to performing multiple particle tracking.

### Immunofluorescence staining

Slices were stained for B and T cell zones as previous described^30^. Briefly, lymph nodes were blocked with purified anti-mouse CD16/32 (BioLegend) and then stained for B cell or T cell zones using primary and secondary antibodies: anti-mouse CD45R/B220-AF647 (BioLegend), anti-collagen III (Proteintech), and donkey anti-rabbit 488 (BioLegend).

### Nanoparticle formulation

500 nm polystyrene beads were densely coated in polyethylene glycol (PEG) using ethyl-3-(3-dimethylaminopropyl) carbodiimide (EDC) and *N*-hydroxysuccinimide (NHS) chemistry, as previously described^60^. Particles were diluted in deionized (DI) water at a 1:4 ratio and sonicated. 5 kDa or 40 kDa amine-terminated PEG (Creative PEGworks) were conjugated to the surface of 500 nm and 1 µm carboxylate-modified fluorescent polystyrene beads (Fluospheres^TM^, ThermoFisher), respectively, using 0.02 mM EDC (Invitrogen) and 14 mM NHS (Sigma) dissolved in 200mM borate buffer (pH = 8.2). The reaction was allowed to proceed on a rotary shaker for at least 4 hours at room temperature. For maximum PEG coverage on 500 nm and 1 µm beads, 1.575 µmol of 5kDa PEG and 0.2175 µmol of 40kDa PEG were used, respectively. Particles were centrifuged for 10 minutes at 15000 x g and 12 minutes at 10000 x g for 500 nm and 1 µm particles, respectively, and washed 3 times using deionized (DI) water. Particles were resuspended at 1% w/v in DI water and stored at 4°C. Particle size and ζ-potential was characterized using dynamic light scattering (DLS) and phase analysis light scattering (PALS) (Brookhaven Instruments).

### Multiple particle tracking (MPT) analysis

Up to 2 µL of a 1:1000 nanoparticle dilution was pipetted on top of each lymph node slice. A coverslip was placed on top of the slice. A ZEISS Axioscope 5 microscope with 63X objective using ZEISS software (ZEN lite) was used to capture videos at a temporal resolution of 30 ms for 10 seconds. Four slices were collected from each node, with 3 videos collected from each slice (representing the B cell zone, T cell zone, and a random zone). Averages for the ‘overall node’ are computed from data collected from all 4 slices across all zones (total of 12 videos per lymph node). Averages for the B and T cell zones ignore all videos collected in the random zone. Ensemble averaged mean square displacement (MSD) was calculated from nanoparticle trajectory using MATLAB^61^ with at least 50 particles per video. MSD was calculated using Δ*r*^2^(*τ*) = [*x* (*t* + *τ*) − *x*(*t*)]^2^ + [*y* (*t* + *τ*) − *y*(*t*)]^2^. The MSD values were then used to extrapolate the microrheological properties of the tissue using the generalized Stokes-Einstein relation^33^, defined as 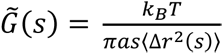, where *k_B_* T is the thermal energy, *a* is the particle radius, and *s* is the complex Laplace frequency. Using the frequency-dependent, complex modulus equation, *G*^∗^(*ω*) = *G*′(*ω*) + *iG*^”^(*ω*), where *s* is substituted with *iω*, we can solve for the elastic and loss moduli. The complex microviscosity (*η*^∗^) can be calculated as *η*^∗^ = *G*^∗^(*ω*)/*ω*, where *ω* is the frequency. Comparisons of loss and elastic moduli were made at a value of 1Hz, in line with standard rheological measurements. The pore size (ξ) is estimated from the MSD^62^ using the equation 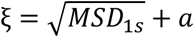, where *a* is the particle radius. Pore size data represent all pore sizes collected from all particles in each slice.

### Gap analysis

Lymph nodes were embedded in optimal cutting temperature compound (Tissue-Tek O.C.T. Compound) and flash frozen via liquid nitrogen. Tissues were stored at −80°C until slicing. 10 *μ*m thick slices were collected using a Leica CM1950 Cryostat. Tissues were then washed with 1x PBS to remove excess OCT and fixed with 2% paraformaldehyde (PFA) in 1x PBS for 5-15 minutes. Tissues were blocked with 0.5% w/v casein in 1x PBS. Slices were imaged using a FV3000 Laser Scanning Confocal Microscope (Olympus) with the following settings: 1.5x zoom with 1024×1024 pixel resolution. The tiling feature was used to create a stitched 3×3 image of the tissue. Z-stack images were captured with 10 slices, each separated by a 0.5-step interval. Images were analyzed using FIJI. The individual stacks were merged using the z-project function at standard deviation projection. Images were then processed using the Gap analysis script^7^. Briefly, images were resized by a factor of 0.4. They were then converted into binary based on a pixel intensity threshold of 0.001. The largest circles possible were fitted into the gaps so that the circles were not overlapping with each other. Circle radii were collected in pixels and converted to *μ*m based on the microscope resolution using the Gap analysis script.

### Statistics

All statistical analyses were performed using GraphPad Prism 10. P values ≤ 0.05 are considered significant. Sample sizes were predetermined based on power analysis (p=0.05, ɑ = 0.8) and guided by prior experience with similar experiments. In general, two-group comparisons were performed using two-tailed, unpaired Student’s t-tests or their non-parametric equivalents depending on data distribution. For multiple group comparisons, one-way ANOVA with Tukey’s multiple comparison test or a non-parametric Kruskal-Wallis test followed by Dunn’s multiple comparisons test were conducted. Normality and homoscedasticity were evaluated prior to analysis to confirm the assumptions of the statistical tests applied. Figure legends include details of the statistical analyses performed for the figure dataset.

## ACKNOWLEDGEMENTS

This work was supported by financial support from NIGMS MIRA 1R35GM142835-01 (KM, VS) and NIH HPI T32 (AR). All animal procedures were performed in accordance with the Guide for the Care and Use of Laboratory Animals and the American Veterinary Medical Association (AVMA) Guidelines for the Euthanasia of Animals and approved by the Institutional Animal Care and Use Committee at the University of Maryland, College Park. We would like to thank Dr. Giuliano Scarcelli, Sachin Suresh, and Anoushka Dasgupta for their inputs on the analysis.

## LIST OF ABBREVIATIONS

ECM: Extracellular Matrix
FRC: Fibroblastic Reticular Cell
MPT: Multiple Particle Tracking
DC: Dendritic Cell
LPS: Lipopolysaccharide
MFI: Mean Fluorescence Intensity
TLR: Toll-like Receptor
PEG: Polyethylene glycol
DI: Deionized
EDC: Ethyl-3-(3-dimethylaminopropyl) carbodiimide
NHS: *N*-hydroxysuccinimide
DLS: Dynamic Light Scattering
PALS: Phase Analysis Light Scattering
MSD: Mean Square Displacement
OCT: Optimal Cutting Temperature
PFA: Paraformaldehyde
PBS: Phosphate Buffered Saline

