## Supplemental Figures for "Nanoparticles reveal permanent and reversible changes to lymph node biomechanics during inflammatory response"

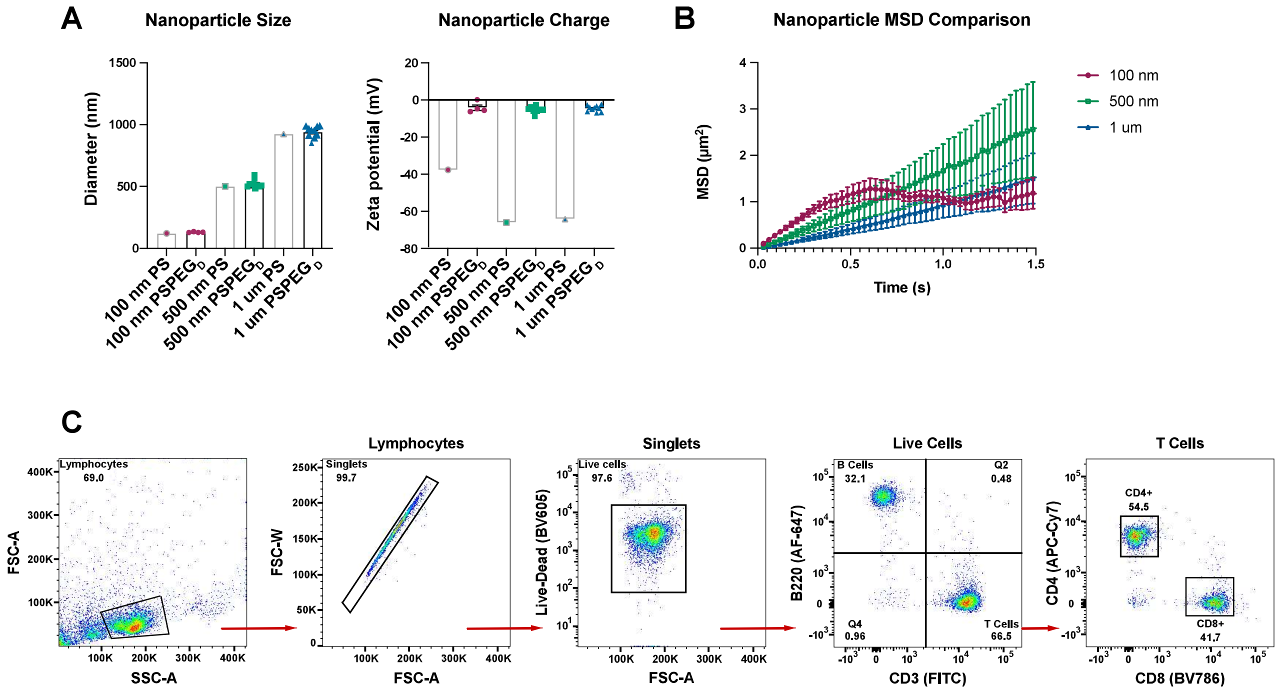


**Supplementary Figure S1** *Nanoparticle characterization of PEGylated polystyrene beads* (A) Size and charge analyzed by dynamic light scattering (DLS) and phase analysis light scattering (PALS). (B) Mean square displacement of $PSPEG_D$ 100nm, 500nm and 1μm polystyrene beads. *Flow cytometry* (C) Dot plot with gating strategy. All values shown in **A** and **B** are mean ± SEM.

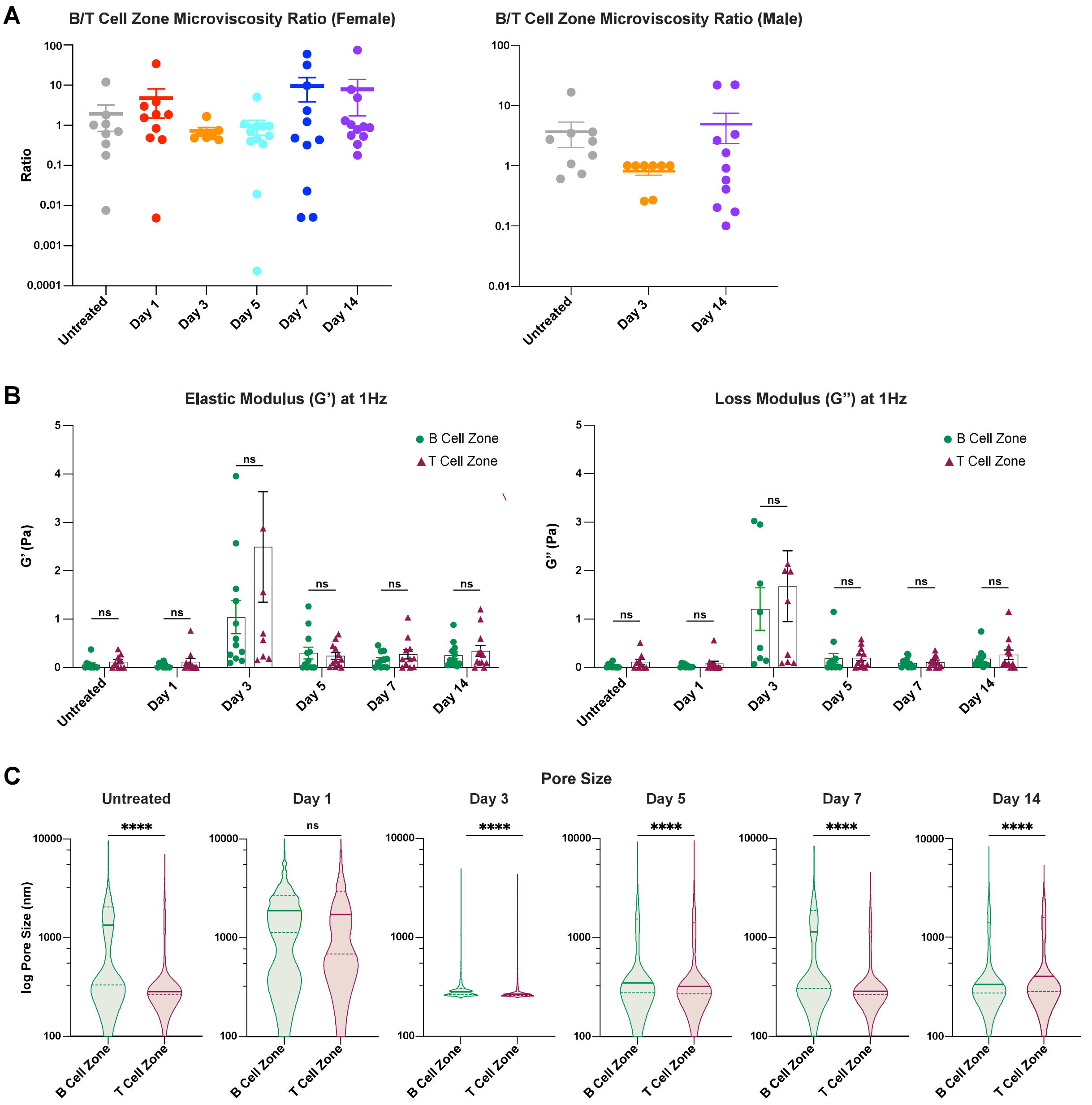


**Supplementary Figure S2** *Comparison of B and T cell zone biomechanics across inflammation* (A) Ratio of microviscosity in B to T cell zones for individual mice in untreated lymph nodes and lymph nodes days 1, 3, 5, 7 and 14 after LPS treatment. (B) Elastic and loss moduli comparison of B and T cell zones at 1Hz. (C) Pore size comparison (by MPT) of B and T cell zones during the course of inflammation. Median and quartile values shown for pore sizes. Other values are reported as mean ± SEM. Y-axis in **C** shown on a logarithmic scale, axis labeled with 'log' to enhance readability. Zone-wise comparison of elastic and loss moduli are done at 1Hz as an average across all mice, and statistical analysis was performed using a Mann-Whitney test on each day **(B).** Statistical analysis for pore size was performed using a Mann-Whitney test on each day **(C).** *p<0.05, **p<0.01, ***p<0.001, ****p<0.0001, ns p≥0.05. N = 10-15 female mice, 8-12 male mice.

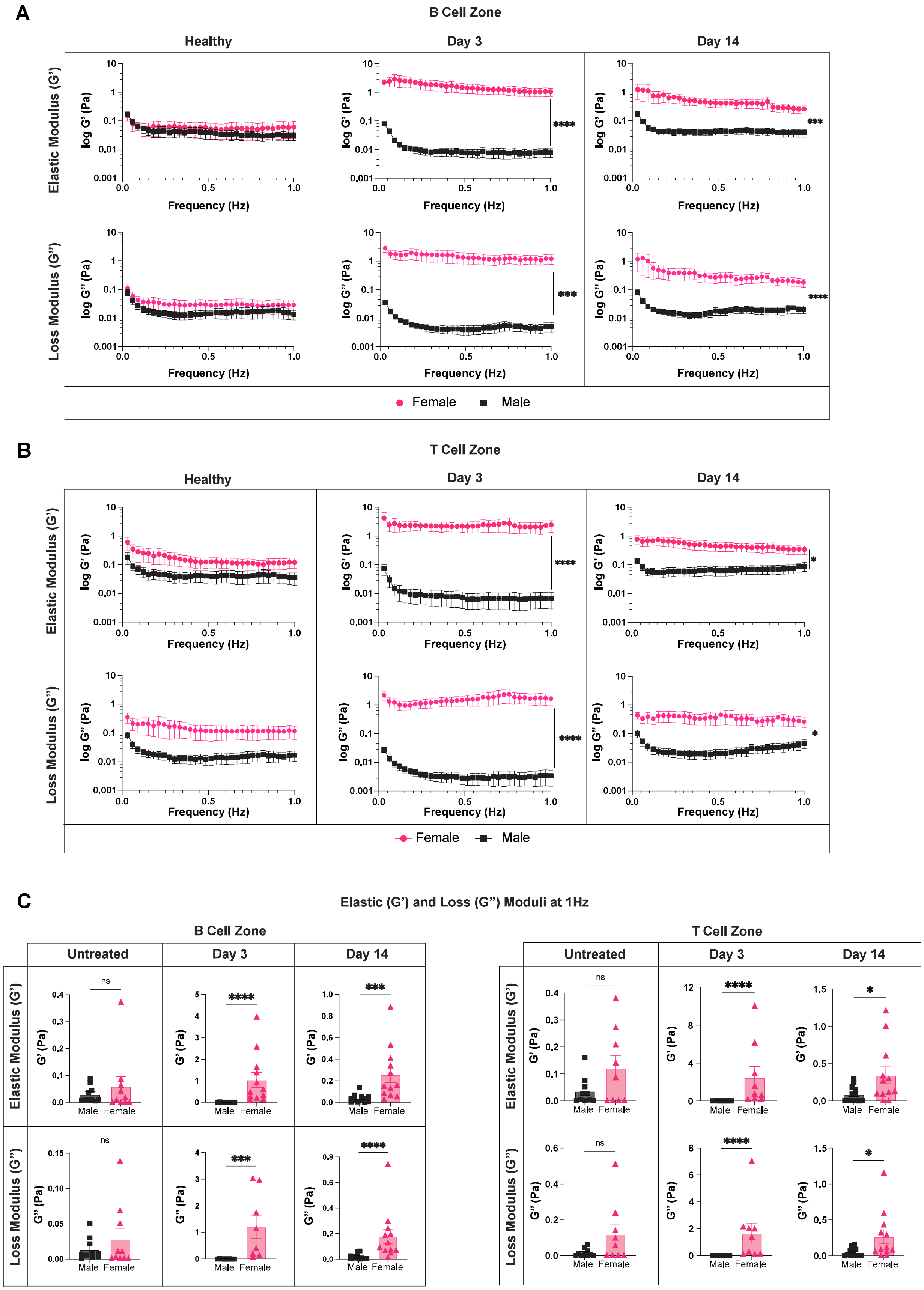


**Supplementary Figure S3**. *Sex differences within B and T cell zones of murine male lymph nodes.* Elastic and loss moduli over 1Hz within the (A) B cell zone and (B) T cell zone. (C) Elastic and loss moduli at 1Hz within B and T cell zones. All values are mean ± SEM. Y-axis in **A,B** shown on a logarithmic scale, axis labelled with 'log' to enhance readability. Elastic and loss moduli are compared at 1Hz as an average across all mice, and statistical analysis is done by Mann-Whitney test(**A-C**). Significance values are denoted on **A,B** for representation. *$p<0.05$, **$p<0.01$, ***$p<0.001$, ****$p<0.0001$, ns $p\geq0.05$. N=8-12 male and female mice.

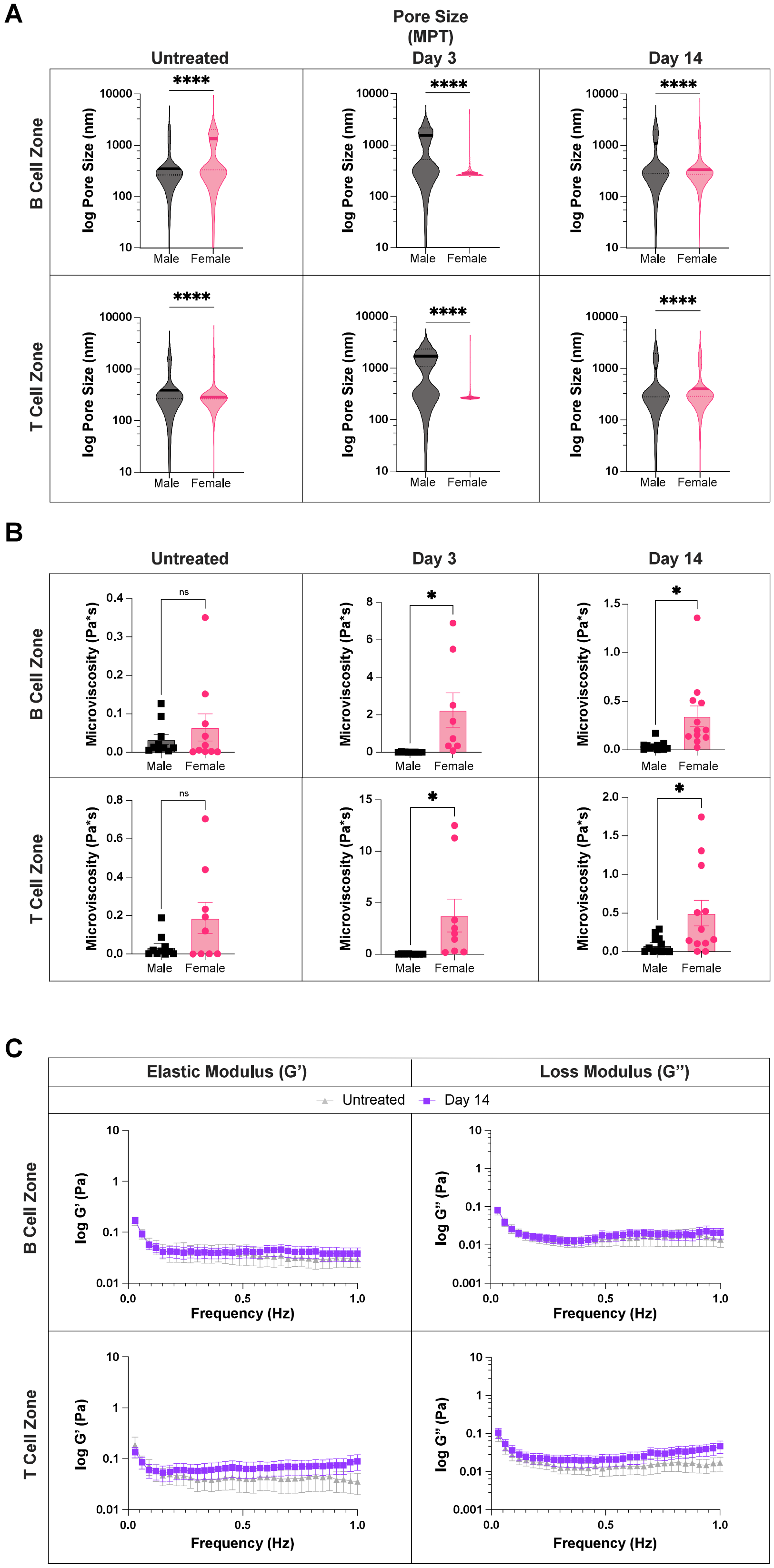


**Supplementary Figure S4** *Sex differences within B and T cell zones* (A) Pore sizes(by MPT) in B and T cell zones (B) Microviscosity in B and T cell zones (C) Elastic and loss moduli over 1Hz in male untreated and day 14 lymph nodes. All values are mean ± SEM. Y-axis in **A,C** shown on a logarithmic scale, axis labelled with 'log' to enhance readability. Statistical analyses of microviscosity data done by Mann-Whitney test **(A).** Elastic and loss moduli are compared at 1Hz as an average across all mice, and statistical analysis is done by Mann-Whitney test **(C).** *p<0.05, **p<0.01, ***p<0.001, ****p<0.0001, ns p≥0.05. N=8-12 male and female mice.

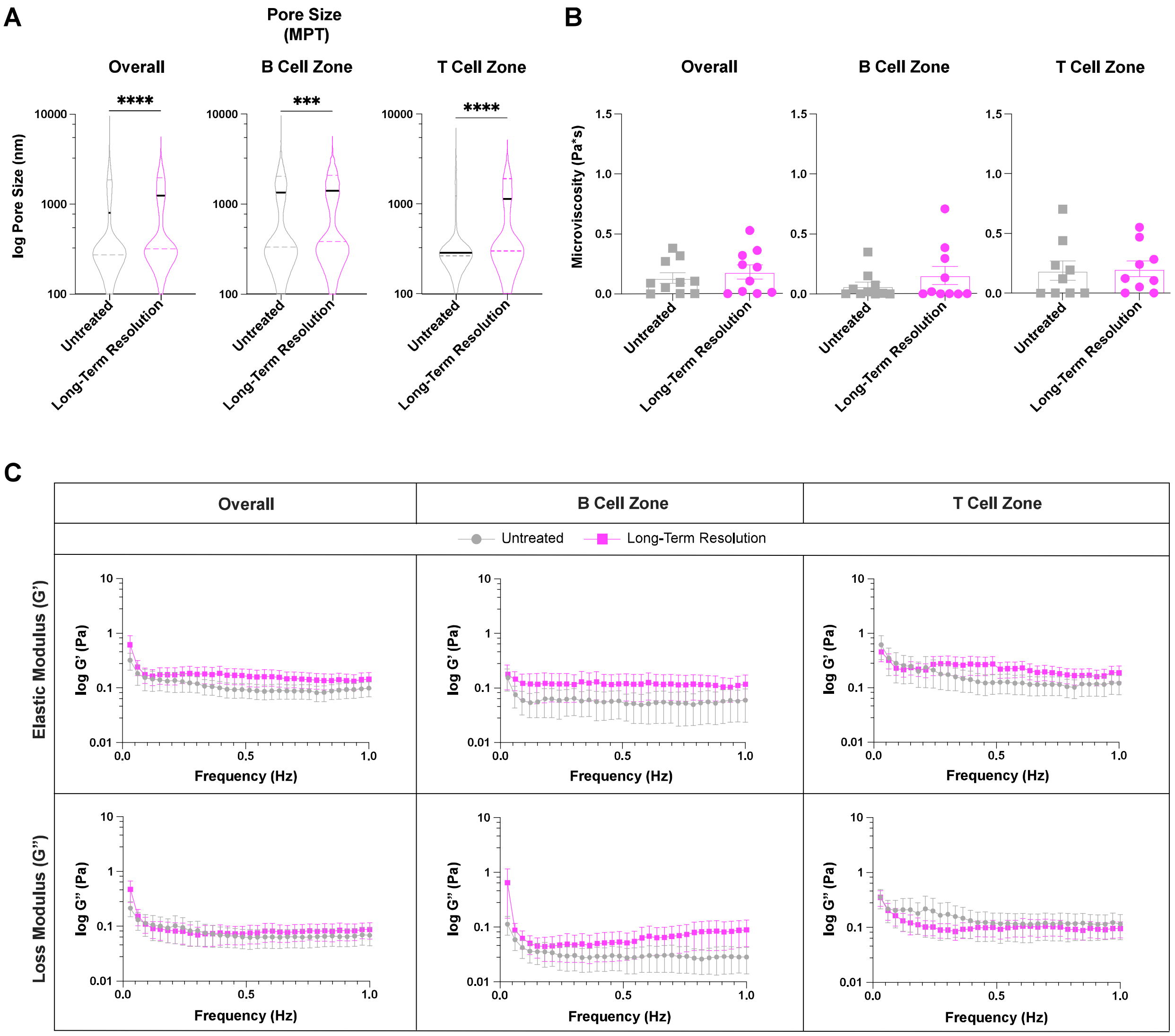


**Supplementary Figure S5** *Chronically inflamed lymph nodes left to recover for 4 weeks, exhibit similar biomechanics to untreated lymph nodes* (A) Pore sizes (by MPT) in the overall lymph node and in the B and T cell zones and (B) in the overall lymph node and in the B and T cell zones (C) Elastic and loss moduli over 1Hz in untreated and lymph nodes after long-term resolution. All values are mean ± SEM. Y-axis in **A,C** shown on a logarithmic scale, axis labelled with 'log' to enhance readability. Statistical analyses of microviscosity data done by Mann-Whitney test **(A,B).** Elastic and loss moduli are compared at 1Hz as an average across all mice, and statistical analysis is done by Mann-Whitney test **(C).** *p<0.05, **p<0.01, ***p<0.001, ****p<0.0001, ns p≥0.05. N=10-15 female mice.
